# PRDM9 forms an active trimer mediated by its repetitive zinc finger array

**DOI:** 10.1101/498428

**Authors:** Theresa Schwarz, Yasmin Striedner, Karin Haase, Jasmin Kemptner, Nicole Zeppezauer, Philipp Hermann, Irene Tiemann-Boege

## Abstract

PRDM9 has been identified as a meiosis-specific protein that plays a key role in determining the location of meiotic recombination hotspots. Although it is well-established that PRDM9 is a trans-acting factor directing the double strand break machinery necessary for recombination to its DNA binding site, the details of PRDM9 binding and complex formation are not well known. It has been suggested in several instances that PRDM9 acts as a multimer *in vivo*; however, there is little understanding about the protein stoichiometry or the components inducing PRDM9 multimerization. In this work, we used *in vitro* binding studies and mass spectrometry to characterize the size of the PRDM9 multimer within the active DNA-protein complex of two different murine PRDM9 alleles, PRDM9^Cst^ and PRDM9^Dom2^. For this purpose, we developed a strategy to infer the molecular weight of the PRDM9-DNA complex from native gel electrophoresis based on gel shift assays (EMSAs). Our results show that PRDM9 binds as a trimer with the DNA. This multimerization is catalysed by the long ZnF array (ZnF) at the C-terminus of the protein and 11, 10, 7 or 5 ZnFs are already sufficient to form a functional trimer. Finally, we also show that only one ZnF-array within the PRDM9 trimer actively binds to the DNA, while the remaining two ZnF-arrays likely maintain the multimer by ZnF-ZnF interactions. Our results have important implications in terms of PRDM9 dosage, which determines the number of active hotspots in meiotic cells, and contribute to elucidate the molecular interactions of PRDM9 with other components of the meiotic initiation machinery.

## INTRODUCTION

In most mammals, including humans and mice, the meiosis-specific protein PR-domain containing protein 9 (PRDM9) was identified to play a key role in regulating and determining the location of recombination hotspots^1–5^. PRDM9 is a multi-domain protein expressed in prophase I in ovaries and testis^6, 7^ that recognizes DNA target motifs and directs double strand breaks (DSBs) to these target sites. Four functional domains have been described for the PRDM9 protein: KRAB, SSXRD, PR/SET, and the C-terminal zinc-finger (ZnF) array. The ZnF array recognizes and contacts specific target DNA sequences^1,8–13^ commonly found at hotspot centres in humans and mice^2, 3^. This ZnF-DNA interaction is very stable and lasts for many hours, which is important for other PRDM9 domains to carry out their activity throughout the different stages of meiotic prophase I and direct the placement of DSBs in leptotene^12^. The PR/SET domain has trimethylation activity and labels surrounding nucleosomes by H3K4me3 and H3K36me3 marks^7,14–17^. The role of H3K4me3 and H3K36me3 in meiosis is not yet fully understood, but these epigenetic marks were shown to co-occur at hotspot regions (^16^ and reviewed in^18^) and are functionally important in the interaction with components of the DSB machinery, located on the chromatin axis^19, 20^. In addition, H3K4me3 is associated with an open chromatin structure at DSB targets hypothesized to be important for proper DNA pairing between homologues and recognition, which would be otherwise hidden within nucleosomes^21^. Finally, the N-terminal KRAB domain (together possibly with the SSXRD domain) binds to other protein complexes, like EWSR1, CDYL, EHMT2^20^ and CXXC1^19, 20^, involved in tethering the target DNA in the loop with the axis where proteins of the DSB machinery are located^22^. Note that PRDM9 interacts with CXXC1, but it is not an essential link for meiotic recombination progression in mice^23^. All four domains of PRDM9 play an important role in the placement of DSBs at hotspot targets recognized by the ZnF-array. Over evolutionary time, species have either lost the complete full length *Prdm9* gene, are missing one of the four domains, or have non-functional changes. In those species lacking a functional PRDM9, DSBs occur at PRDM9-independent sites such as transcription start sites (TSS) or CpG islands, as observed in birds and dogs^24–28^.

PRDM9 has been shown to be highly polymorphic between and within species. Most mammals, like humans^3–5,29^, chimps^30–32^, mice^1,4,10,33^, equids^34^ and cattle^35^ harbour different *Prdm9* alleles, which have a change in amino acids contacting the DNA, and vary in the arrangement and number of ZnFs within the DNA-binding domain. This results in the activation of different sets of hotspots. In heterozygous individuals, the *Prdm9* diversity can affect hotspot activation, since different alleles do not show additive behaviour but rather compete for DNA binding. This results in predominance of hotspots from one allele and suppression of hotspots targeted by the other allele, as it was shown in both humans for the C vs. A allele^5^ and mice for the 9R vs. 13R allele^2^ or the B6 vs. CAST allele^33, 36^.

*Prdm9* has been identified also as a speciation gene playing an essential role in hybrid sterility^37^. Interestingly, only certain combinations of heterozygous *Prdm9* alleles are incompatible in a specific genetic background^38, 39^. The process is not yet fully understood, but it has been observed that the heterozygous *Prdm9* alleles preferentially activate target sequences at the non-self homolog, which is influenced by sequence erosion at recombination hotspots. This leads to an asymmetric binding of PRDM9, and thus also an asymmetric distribution of double strand breaks (DSB) between homologues, which is linked with chromosome asynapsis and hybrid sterility(^33, 40^ and reviewed in^21^). Moreover, it has been shown that PRDM9 dosage also determines the number and activity of hotspots. Hemizygous null mice (*Prdm9^+/-^*) with only one copy of *Prdm9,* have a fewer number, and less active recombination hotspots resulting in reduced fertility^41^. Complete loss of *Prdm9* leads to sterility in both sexes, since initiated DSBs are not properly repaired, which causes asynapsis and disrupted gametogenesis at the pachytene stage resulting in meiotic arrest^7^. Moreover, certain heterozygous F1 hybrids also show partial asynapsis with a strong bias towards the smallest autosomes, as it was shown for PWD x C57BL/6 crosses (*Prdm9^Msc/Dom2^*), which could be rescued by introducing a minimum of 27 Mb consubspecific homologous sequence to one of the chromosomes pairs restoring the symmetric hotspot distribution^42^. It was also demonstrated that removing or overexpressing a certain PRDM9 allele, and therefore increasing the PRDM9 dosage, could rescue fertility in sterile hybrid crosses^39^. This suggests that a certain number of active hotspot sites are required for successful meiotic progression, which among others is controlled by the dosage of PRDM9.

Recently, it has been observed that PRDM9 can form functional multimeric complexes^14, 36^. How this multimerization affects the activity of PRDM9 is not known, but it could play a role in the preferential hotspot usage by the dominant PRDM9 allele in heterozygous individuals, where the binding is driven by the stronger allele^36^. Multimerization could also play a role in hybrid sterility, in which otherwise active alleles are sequestered in a heteromer resulting in a more symmetric distribution of DSBs and synapsis (reviewed in^21^).

To date observations of PRDM9 multimerization are based on cell systems over-expressing two different alleles of PRDM9 with distinct tags^14, 36^. However, it is not known how many PRDM9 units form the multimer, which is key information to understand how PRDM9 interacts at a molecular level and also influences PRDM9 dosage, especially in the context of PRDM9 heterozygosity. In addition, it is still unknown whether different ZnF arrays within a multimeric complex interact also with multiple DNA targets. By addressing these aspects, we will gain important insights in the nature of the ZnF-DNA interaction and DNA targeting.

In this work, we performed an *in vitro* analysis of the DNA-PRDM9 complex using electrophoretic mobility shift assays (EMSA) to infer the stoichiometry of the active complex. We show that the molecular weight (MW) of a complex can be inferred from its electrophoretic migration distance under non-denaturing conditions. In combination with mass spectrometry, we estimated that PRDM9 forms a trimer when actively bound to DNA. This trimer was observed for two different PRDM9 alleles, PRDM9^Cst^ and PRDM9^Dom2^. Moreover, the trimer formation is mediated within the variable ZnF array and at least 5 out of 11 ZnFs are sufficient to form a stable DNA-binding trimer. Finally, our data suggest a model in which only one of the ZnF array is involved in DNA binding; whereas, the other two ZnFs likely are involved in protein-protein interactions.

## RESULTS

### Uncoupled binding of PRDM9 with linked successive target sequences

In order to better understand the binding behaviour of PRDM9 to its target DNA, we first investigated with *in vitro* gel shift assays how the ZnF complex interacts with its DNA target. Can the different PRDM9 units in a multimer interact simultaneously with several DNA binding sites? If so, is there a cooperative binding to the DNA between the units in the multimer? We answered these questions with EMSA, a standard molecular biology technique based on native gel electrophoresis used to analyse a DNA-protein complex visualized by its slower migration compared to free DNA. We designed DNA fragments with one (single-Hlx1) or two adjacent target sites (tandem-Hlx1) derived from the *Hlx1* hotspot known to specifically bind the PRDM9^Cst^ ZnF array (ZnF^Cst^) of *Mus musculus castaneus* origin^9,12^. If different units (single PRDM9 proteins) within the PRDM9 multimer interact simultaneously with the two DNA binding sites of the tandem-Hlx1, then there is no change in the overall MW of the complex and migration distance. In contrast, if two independent complexes form with the tandem-Hlx1, then this higher MW complex has a slower migration.

For the design of the single and tandem-Hlx1, we considered previous experiments showing that ZnF^Cst^ bound specifically 34 nucleotides, yet unspecific flanking DNA improved the binding^12^. Thus, the DNA sequence contained either one or two adjacent 34bp specific target sites plus 20-23bp flanking regions (single-Hlx1 or tandem-Hlx1 with 75bp or 114bp, respectively), as shown in Figure S1. We analysed the binding of these two DNA fragments to different protein concentrations of ZnF^Cst^ coupled to a maltose binding domain (hereafter MBP-ZnF^Cst^) in an EMSA titration experiment.

We observed that the single-Hlx1 DNA formed a complex (shifted band) at very low protein concentrations. The intensity of the shift increased with protein concentration saturating the free DNA and forming a complex (Figure 1A), as observed before^12^. For the tandem-Hlx1, two different states of the complex were detected with increasing protein concentrations: a lower-shift and a super-shift. The lower-shift was observed at low PRDM9 concentrations, and as PRDM9 concentrations were increased, a second super-shift became visible (Figure 1B). A super-shift is observed regularly in EMSA when a second protein (e.g. antibody against the protein) is incubated with the complex, resulting in a large change of the overall MW slowing the migration of the complex to a super-shift^43^. In our case, the super-shift can be explained by the binding of an additional PRDM9 to the second available *Hlx1* site with increasing protein concentrations. Since the binding sites are on one DNA strand, the two complexes stay linked forming a super-shift. The dynamics are as follows: the lower-shift increases at low PRDM9 concentrations until half of the sites are filled (Figure 1B and 1D). With further increase in protein, the second *Hlx1* target site gets bound, with the effect that the intensity of the lower-shift diminishes and is replaced by an increasing super-shift. The overall free DNA decays at the same rate for both the single and tandem-Hlx1 (Figure 1A-1D). These results demonstrate that the multiple ZnF arrays within a multimer do not interact simultaneously with several DNA binding sites. Instead, the second free site on the tandem-Hlx1 is bound by the ZnF array of an independent PRDM9 multimer. An alternative explanation is that PRDM9 is a monomer, and thus only one ZnF array interacts with the DNA; however, this is not the case as we show in the next sections.

**Figure 1.**
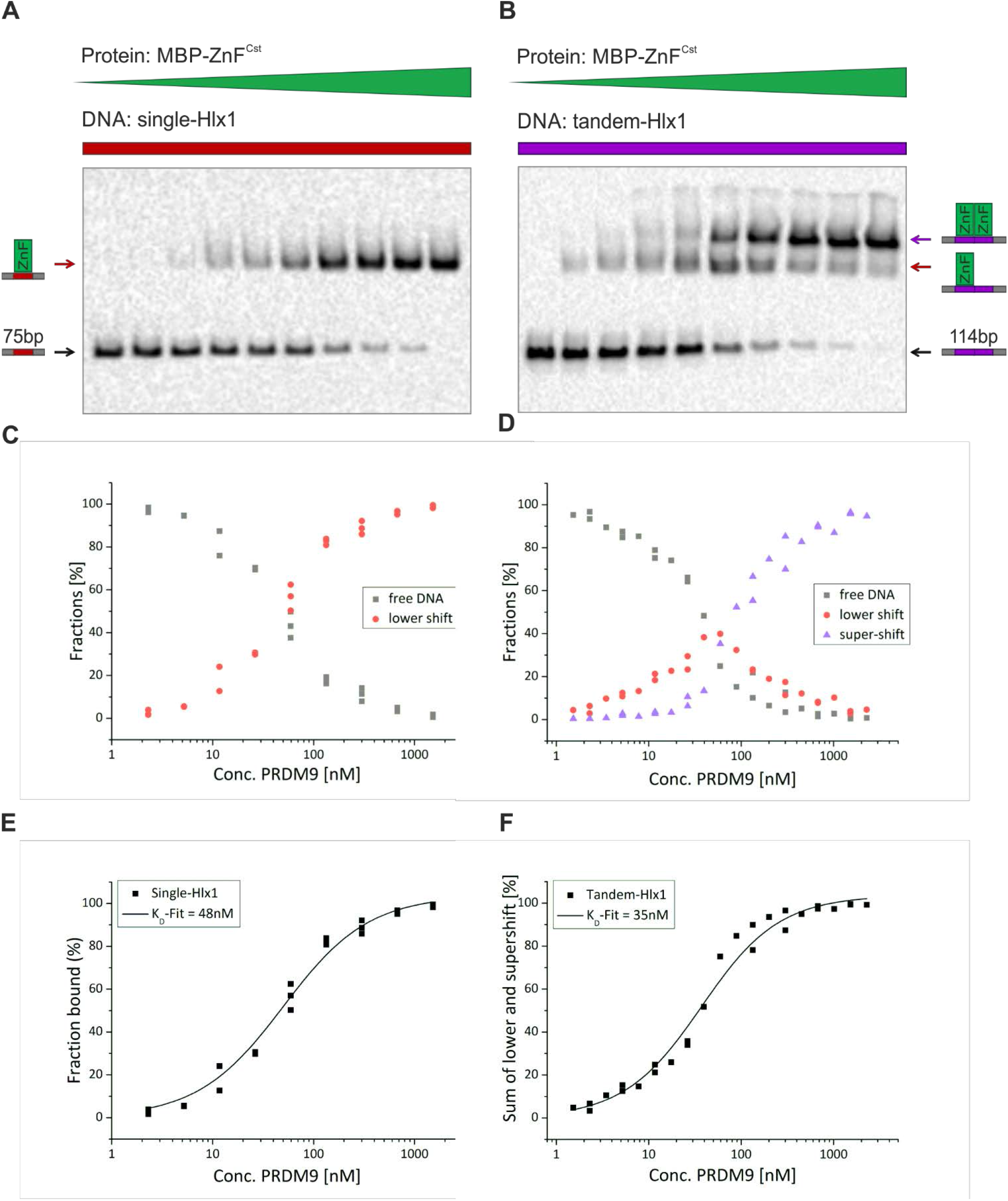
Binding of the PRDM9-ZnF to one or two consecutive target sites. **(A+B)** Shown are titration EMSA experiments in which serial dilutions of MBP-ZnF^Cst^ (2.5μM – 1.5nM) were incubated with constant amounts of labelled target DNA (5nM). Two different DNA targets were used, (A) single-Hlx1 with a length of 75bp and (B) tandem-Hlx1 with a length of 114bp, the latter carrying two consecutive *Hlx1* binding sites. The lowest band (black arrow) is the unbound DNA and the shifted bands are the complex with of either one (red arrow) or two (purple arrow) proteins at the target DNA, labelled as lower-shift and super-shift, respectively. Pixel intensities of the unbound and shifted bands were quantified using the ImageLab software (BioRad). **(C+D)** Different fractions (% fraction) of the binding reaction (fraction unbound = free DNA, grey; fraction after complex formation containing 1xPRDM9, red; and the super-shift fraction containing 2xPRDM9, purple) were plotted against the PRDM9 concentration at a semi-logarithmic scale with the OriginPro8.5 software (OriginLab). **(E+F)** The fraction bound [FB=shift/(shift+unbound)*100] was plotted against the PRDM9 concentration in a semi-logarithmic scale and a K_D_-fit was performed using a function for receptor-ligand binding in solution (as it was described in^12^). The K_D_ for the (E) single-Hlx1 and (F) tandem-Hlx1 (sum of lower- and super-shift) was estimated to be 48nM and 35nM, respectively.

A quantitative analysis (Figure 1E and 1F) shows that the intensity of the sum of the shifts (lower + super-shift) is correlated directly with the affinity of the ZnF. We estimated that the tandem-Hlx1 DNA has a similar affinity to the ZnF as the single-Hlx1 (K_D_ = 35nM and 48nM, respectively), also corroborating that the tandem-Hlx1 is bound by two independent PRDM9s. Moreover, the similar affinity constants between the single- and tandem-Hlx1 also indicate that there is no cooperativity effect between the binding of different complexes on adjacent PRDM9 specific sequences. Note that these K_D_ values are slightly higher than obtained with the same approach in a previous work (24.5nM ± 2.6)^12^). A possible reason for this deviation in the K_D_ could be the much shorter incubation times used here (60 minutes vs. 90 hours) with an effect in the equilibrium states and ultimately the K_D_ when loading the EMSA.

### The molecular weight of the PRDM9-DNA complex can be determined by native gel electrophoresis

Native gel electrophoresis can be used to infer the molecular weight (MW) of negatively charged, linear chains such as DNA or SDS-denatured proteins, for which the MW is inversely proportional to the logarithm of the migration distance in a gel^44–46^. The migration of these linear, negatively charged chains is independent of the total charge and conformation of the molecule and follows the ‘reptation principle’. This model proposes that the negative charge on one end of the molecule is sufficient to drive the rest of the molecule that migrates snakelike through the pores of the gel, oriented by the negative charge on one end and pulling the rest of the molecule through the same path^45–49^ (for more details see Supplementary_Notes).

We developed two different strategies, *assay I* and *assay II,* to infer the molecular weight of the DNA-PRDM9 complex in a polyacrylamide gel under non-denaturing conditions. As before, we used EMSA for visualizing the mobility of the complex and further estimate the protein stoichiometry by comparison to a standard series. In both assays, the migration of the complex was driven by the reptation of the long linear DNA overhangs flanking the complex.

In the more conservative *assay I,* our standards were PRDM9 ZnF complexes (ZnF + DNA) with a constant conformation-charge, but different molecular weights given by the length of the flanking DNA. Previous methods used a similar strategy of constant charge and conformation to derive a function of relative migration distance vs. molecular weight in a Ferguson plot^44,50,51^. For this purpose, we used in *assay I* the tandem-Hlx1 known to form one or two linked complexes, as described in the previous section. Specifically, we designed DNA fragments of different lengths with one binding site (single-Hlx1) or two consecutive (tandem-Hlx1) binding sites, all with increasing non-specific flanking sites (Figure 2A and Figure S1) resulting in a lower-shift (red rectangles) for the single-Hlx1 or lower- and super-shift (purple rectangles in Figure 2B) bands for the tandem-Hlx1 sequences. The lower-shifts (one complex) were used as standards to infer the MW of the second complex in the super-shifts (Figure 2B and Figure S2). The standard curve with nine measurements resulted in a very high correlation of a linear regression function plotted in a log-scale (Figure 2C). The MW of the protein constructs was then estimated from the derived regression function as the average of four independent measurements (super-shift) within one experiment (Figure 2B-C and Table S1).

**Figure 2.**
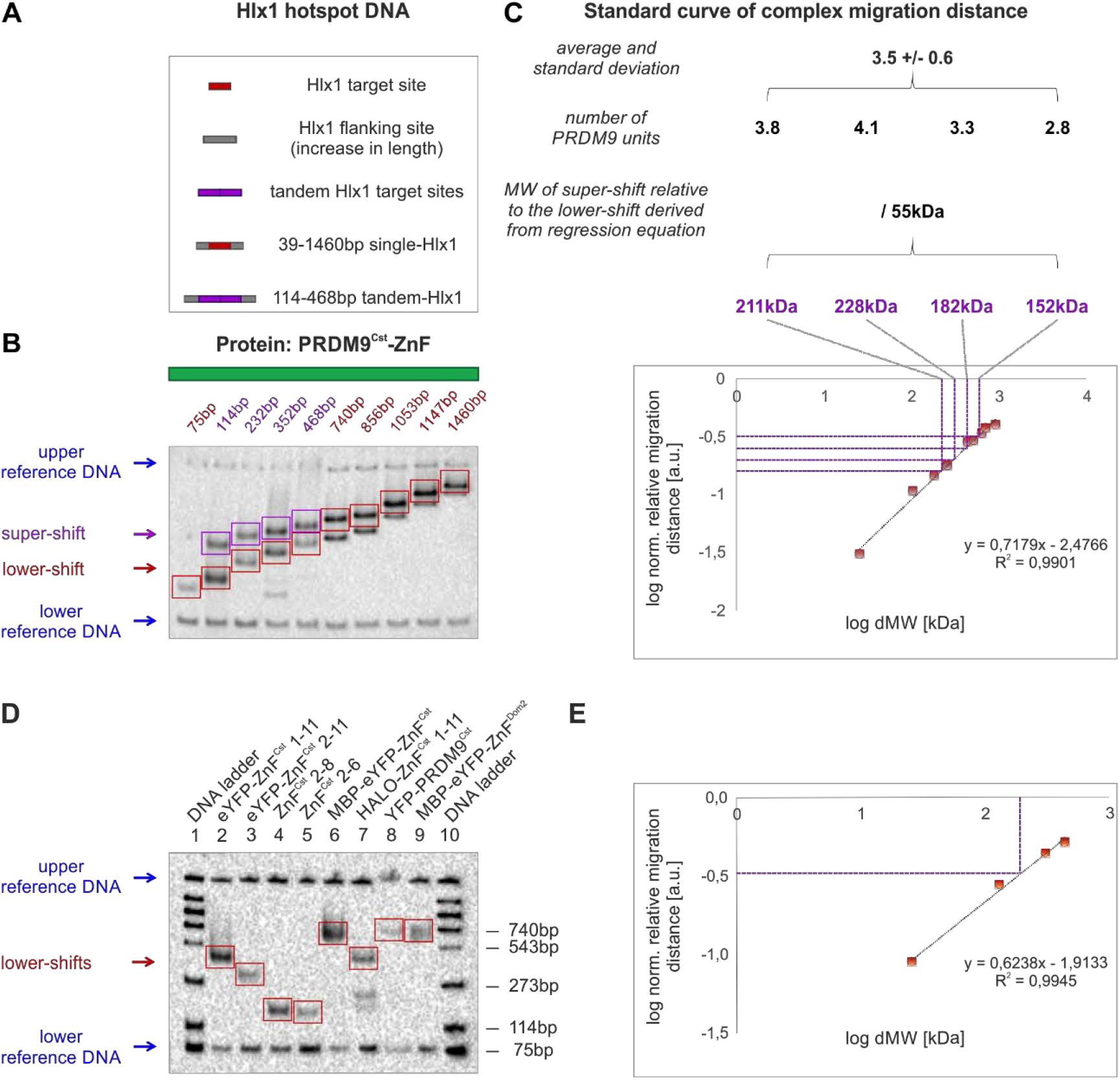
Two strategies to infer the molecular weight of PRDM9 from native gel electrophoresis. **(A)** Different sizes of biotinylated DNA containing one (red) or two (purple) *Hlx1* binding sites (34bp minimal target site for PRDM9^Cst^) were used as DNA standards. The DNA fragments increase in non-specific flanking sites (grey). **(B)** *Assay I:* DNA carrying one or two protein complexes was separated by a native polyacrylamide gel resulting in lower- and super-shift bands (red and purple arrows/rectangles, respectively). Blue arrows indicate long (4368bp) and short (220bp) reference DNA, tested not to interact with the protein, used to normalize for migration distance in each lane. Note that for high MW fragments, the free DNA shows up also on the gel but was not used for the analysis. **(C)** The migration distance of the PRDM9-DNA complexes (lower-shift), relative to the complex in the first lane (75bp single-Hlx1) was plotted against the known relative increase in molecular weight (dMW) between DNA targets in a log-scale. The difference in migration distance of the super-shift relative to the lower-shift of four tandem-Hlx1 fragments was used 1) to estimate the MW representing the second protein complex using the regression equation; 2) to calculate the number of PRDM9 units based on the MW of the PRDM9 construct; and 3) to determine the average and standard deviation of the units from the four tested tandem fragments. Note that complexes with lower molecular weight get resolved better in electrophoresis and the estimation of the molecular weight from the migration distance is more accurate. **(D)** *Assay II:* Binding complexes of eight different PRDM9 constructs with single-Hlx1 75bp for PRDM9^Cst^ constructs and single-Pbx1 75bp for PRDM9^Dom2^ construct (lower shifts, red arrow/rectangles) were separated on the native EMSA gel. Lane 1 and 10 show a DNA ladder, with the respective fragment lengths shown on the right. Each lane included a short (75bp) and long (75bp, loaded 10min before termination of electrophoresis) reference DNA (blue arrows) used to normalize the migration distance within each lane. The measurements were performed in four replicates of independent experiments. **(E)** The normalized migration distance of the DNA ladder bands in lane 1 and 10 relative to the shortest, 75bp, molecule was plotted against the relative increase in MW in a log-scale. The resulting regression equation was used to calculate the MW of the lower-shift complexes and the number of protein units within the complex were estimated as described in panel C.

In the simplified *assay II,* the MW of the different protein constructs was inferred by comparing the migration of the complex directly to free DNA standards (Figure 2D-E). In order to further validate this strategy, we assessed PRDM9 constructs with different charges and conformations by adding different tags and PRDM9 domains, originating from the PRDM9^Cst^ and PRDM9^Dom2^ variants (Figure 3A). By comparing the migration of the shifted bands (lower-shifts, red rectangles) relative to the migration of a DNA ladder (free DNA of different sizes), we estimated the MW of the PRDM9 complex and derived the protein units within each construct (Figure 3B, Figure S3 and Table S1).

**Figure 3.**
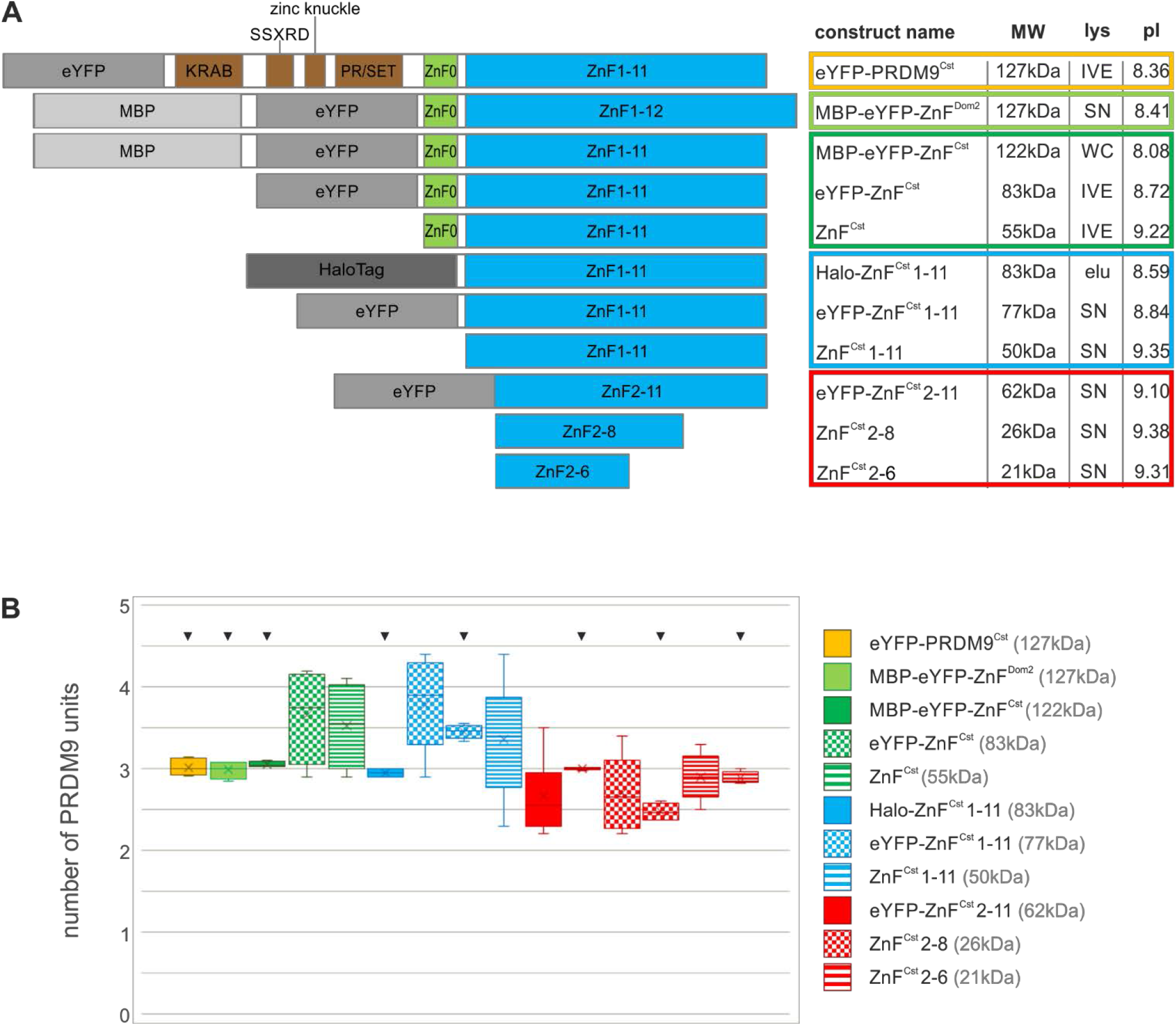
PRDM9 multimerization is mediated within the ZnF array. **(A)** Different PRDM9 constructs used in this study are shown. Domains of PRDM9 are color-coded and additional tags are shaded in grey. Construct name, size, expression system (lys) and theoretical pI are shown on the right in a table format. Cell-free *in vitro* expression, IVE; bacterially expressed whole-cell fraction, WC; bacterially expressed soluble fraction, SN; semi-pure elution via ion exchange chromatography, elu. **(B)** Box plot of the tested PRDM9 constructs representing the distribution of measured PRDM9 units within a multimer complex of assay I and II. Different PRDM9 constructs are color-coded: yellow, full-length PRDM9^Cst^; light green, ZnF domain of PRDM9^Dom2^; dark green, ZnF domain of PRDM9^Cst^; blue, tandem ZnF array of PRDM9^Cst^ without ZnF0; red, truncated ZnF array of PRDM9^Cst^. Markings on top indicate results from assay II to distinguish from results of assay I.

We compared the two developed assays by testing four ZnF^Cst^ constructs with both methods and did not observe differences in the estimated protein stoichiometry (Table 1 and Table S1). This indicates that the migration of the complex in the native gel is driven invariably by the reptation of the long flanking DNA chain, independent of protein charge or conformation.

**Table 1.** Multimerization measured by assay I and II. Four different PRDM9 truncated ZnF constructs measured in both *assay I* and *II* resulted in comparable average estimates of protein stoichiometry. The confidence intervals are given in parenthesis. The size of each construct (MW in kDa), the used expression system (bact. SN, soluble fraction of bacterial expression; WC, whole-cell fraction including cell debris of bacterial expression), and the theoretical isoelectric point (pI) are shown.

| PRDM9 construct | construct name | MW [kDa] | expression system | pI | protein stoichiometry |  |
| --- | --- | --- | --- | --- | --- | --- |
|  |  |  |  |  | assay I | assay II |
| Truncated ZnF array | eYFP-ZnF<sup>Cst</sup> 1-11 | 77 | bact. SN, WC | 8.84 | 3.8 (0.46) | 3.5 (0.09) |
|  | eYFP-ZnF<sup>Cst</sup> 2-11 | 62 | bact. SN | 9.1 | 2.7 (0.44) | 3.0 (0.01) |
|  | ZnF<sup>Cst</sup> 2-8 | 26 | bact. SN | 9.38 | 2.7 (0.46) | 2.5 (0.11) |
|  | ZnF<sup>Cst</sup> 2-6 | 21 | bact. SN | 9.31 | 2.9 (0.30) | 2.9 (0.08) |

### PRDM9 interacts with the DNA as a trimer

In order to assess the protein stoichiometry of the PRDM9 multimer and the PRDM9 domain mediating this multimerization, we designed eleven different protein constructs missing selected domains of the PRDM9^Cst^ and PRDM9^Dom2^ variants (Figure 3A). In addition, constructs carried different tags like eYFP (enhanced yellow fluorescent protein), MBP (maltose binding protein), or Halo (His_6_-HaloTag) and were produced by distinct expression systems, like cell-free *in vitro* expression (IVE) or bacterial expression and protein lysate protocols (Table S1 and Supplementary_Methods). The majority of the bacterially expressed constructs were used as crude lysates without further purification obtained from the whole-cell fraction (WC) with cell debris or from the soluble fraction (SN) excluding cell debris. Only the lysate preparation for the construct containing the Halo tag was semi-pure and included a purification step based on ion exchange chromatography by using SP Sepharose (for details see Supplementary_Methods).

We assessed that the full-length PRDM9 interacts with DNA as a trimer (Figure 3B and Table S1). In fact, all of our tested constructs, including the ZnF domain of two different murine alleles, PRDM9^Cst^ and PRDM9^Dom2^ without the KRAB, SSXRD and PR/SET domains bound as a trimer (MBP-eYFP-ZnF^Dom2^, MBP-eYFP-ZnF^Cst^, eYFP-ZnF^Cst^, ZnF^Cst^). We further removed individual ZnFs from the ZnF domain starting with ZnF0 (spaced 102 amino acids from the tandem array ZnF 1-11) of PRDM9^Cst^ (Halo-ZnF^Cst^ 1-11, eYFP-ZnF^Cst^ 1-11, ZnF^Cst^ 1-11), eYFP-ZnF^Cst^ 2-11 (missing ZnF0 and ZnF1), ZnF^Cst^ 2-8, and ZnF^Cst^ 2-6. Interestingly, even the smallest ZnF2-6 construct (with only five out of eleven ZnFs of PRDM9^Cst^) bound as a trimer with the DNA. This strongly suggests that the trimer formation of active PRDM9 is mediated within the variable DNA-binding ZnF array and at least five out of eleven fingers are sufficient to form a stable DNA-binding multimer. Moreover, we demonstrated that the PRDM9 trimerization is not dependent on the PRDM9 allele, since both PRDM9^Cst^ and PRDM9^Dom2^ did show the same protein stoichiometry.

We analysed the data from *assay I* and *assay II* independently with an ANOVA test and observed some differences between the PRDM9 constructs (detailed analysis can be found in Materials and Methods and Supplementary_Statistical_Analysis). Since, these differences can neither be explained by construct size, additional tags, expression system nor theoretical isoelectric point (Figure 3 and Table S1), we suggest that this is due to experimental variations.

### PRDM9 complex binds only one DNA molecule at a time

Since PRDM9 forms a multimer, we also asked whether the different ZnF arrays within the trimer can interact with more than one DNA molecule. The results of the tandem-Hlx1 experiment described initially suggested that the multiple ZnF arrays within the trimer do not interact simultaneously with more than one DNA binding site. However, it is possible that the simultaneous interaction of multiple ZnF arrays with the two adjacent target sites might have posed a physical constraint by the closely spaced target sites in the Hlx1-tandem sequence. Thus, we performed an additional test to assess if the multiple ZnF arrays within the trimer can interact with more than one DNA molecule.

This time we designed an experiment in which PRDM9 was incubated with a short and a long DNA sequence with the same binding site, but unspecific flanking regions of different sizes. Each DNA-protein complex formed a unique shift in EMSA given the difference in MW of the DNA. In model 1, the trimer binds only one DNA (either the long or the short DNA), and we expect two shifts in addition to the two free DNA sequences (Model 1). Alternatively in model 2, the trimer binds two or more DNA sequences, and we expect five bands: three shifts and two free DNA sequences shown in Figure 4A. Our results clearly show the formation of a DNA-protein complex with either the short or the long DNA, but not both, demonstrating that only one of the three ZnF arrays in the multimer actively binds to the DNA (Figure 4B). The remaining two ZnF arrays could be involved in protein-protein interactions stabilizing the multimer.

**Figure 4.**
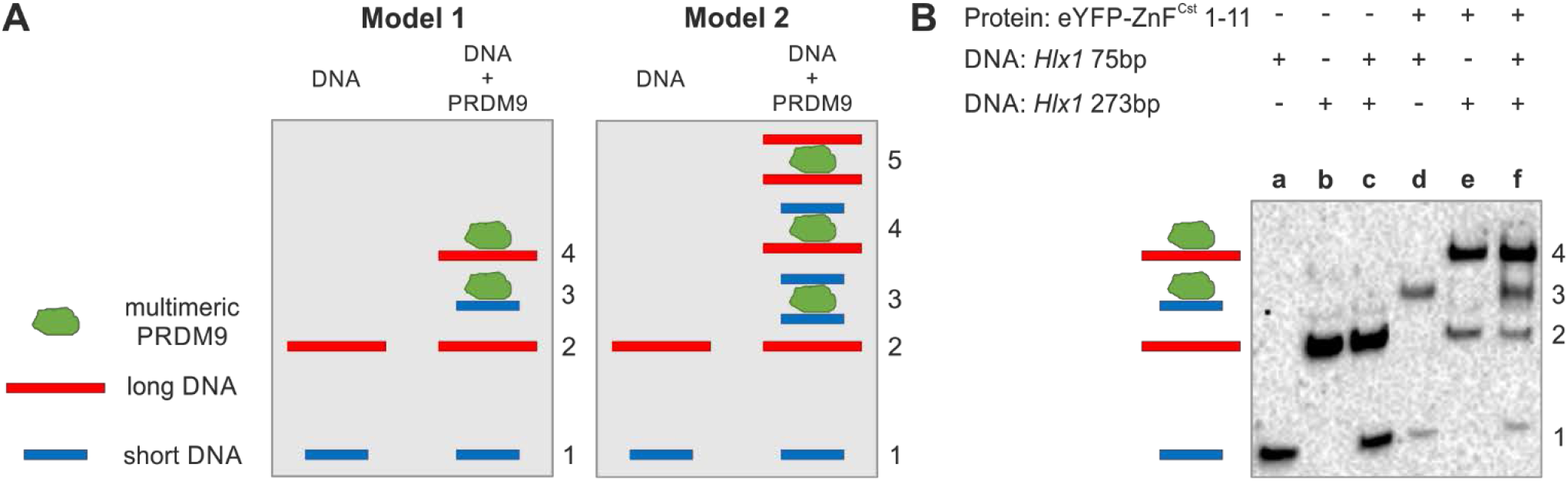
PRDM9 complex forms with only one target molecule. **(A)** The two models represent the binding of the multimeric PRDM9 complex (green) to a short and long DNA (blue and red, respectively) containing the same binding site. The final molecular weight of the protein-DNA complexes varies, resulting in distinct migration distances in the EMSA gel. When mixing equimolar amounts of short and long DNA with PRDM9, the protein will randomly bind either the short or the long DNA. Model 1 represents the banding pattern if the protein complex binds only one DNA molecule resulting in four different bands: (1) short DNA, (2) long DNA, (3) protein + short DNA, (4) protein + long DNA. Model 2 shows the banding pattern if the multimeric protein binds two DNA molecules at a time resulting in 5 different bands:: (1) short DNA, (2) long DNA, (3) protein + 2x short DNA, (4) protein + 1x short DNA + 1x long DNA, (5) protein + 2x long DNA. **(B)** EMSA experiment was performed with eYFP-ZnF^Cst^1-11 and two DNA fragments of the *Hlx1* hotspot DNA that differ in size, 75bp and 273bp mixed at equal molarities (5nM) with 0.25μl eYFP-ZnF^Cst^ 1-11.

### Mass spectrometry demonstrates that the complex is formed by PRDM9 and DNA

So far, our calculations have considered that the PRDM9-DNA complex is mainly formed by these two molecules. Given that our constructs were mainly protein lysates, it is possible that other peptides might be part of the binding complex. In order to test this, we isolated the complex of the semi-pure Halo-ZnF^Cst^ 1-11 protein and the Hlx1-75bp DNA from a native gel (Figure S4A) and analysed it by MALDI-TOF mass spectrometry (Figure 5).

**Figure 5.**
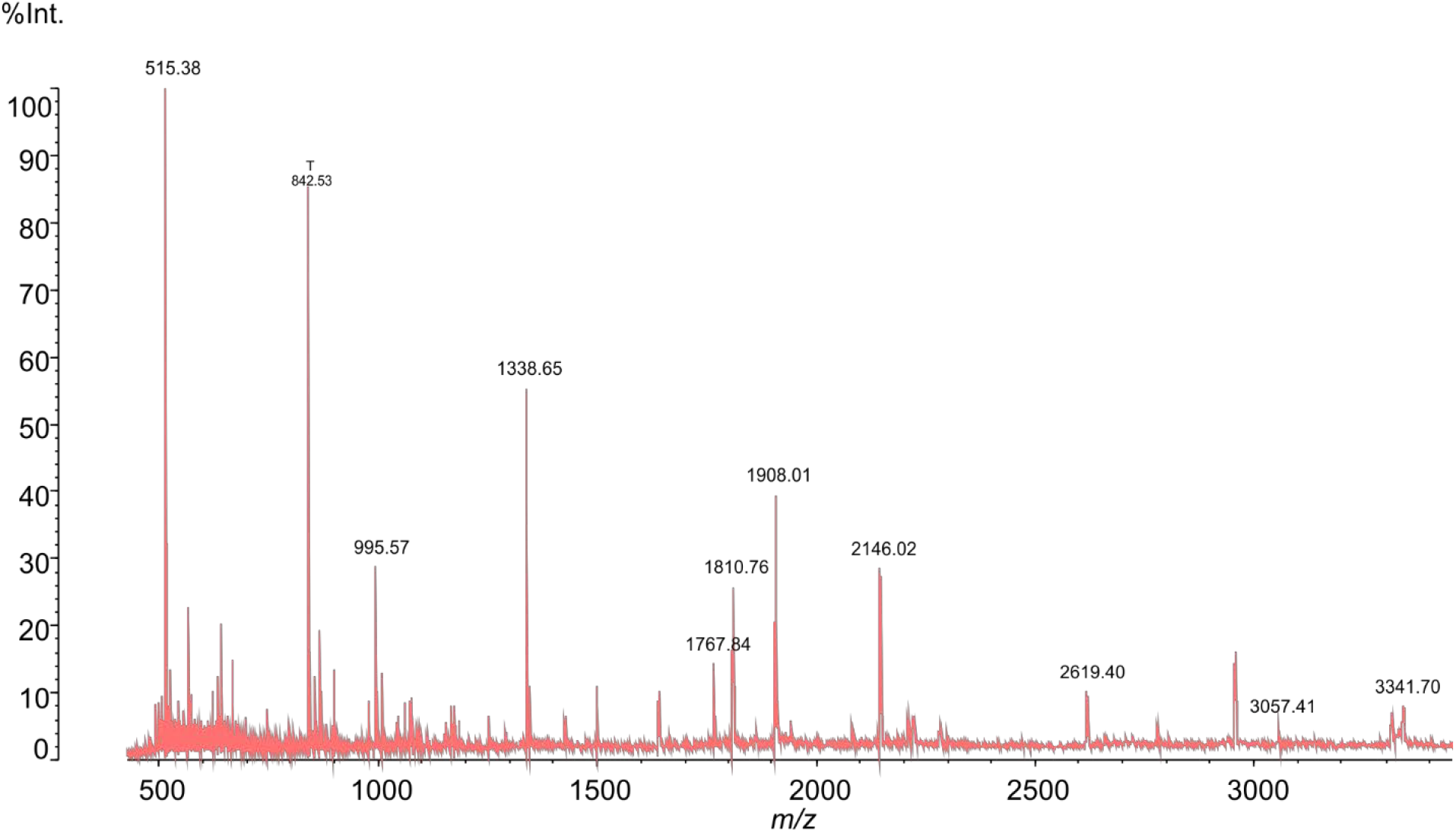
Peptide mass fingerprint spectrum of Halo-ZnF^Cst^ 1-11. Shown is the mass spectrum of Halo-ZnF^Cst^ 1-11, which was incubated with a 75bp DNA fragment from the *Hlx1* hotspot for 60min and cut out from a native 5% acrylamide gel after coomassie staining. After treatment with trypsin digestion, DTT reduction and carbamethylation using iodacetamide, the complex was analysed using a MALDI-TOF Axima Performance instrument (Shimadzu). The spectrum shows the peptide mass fingerprint of the PRDM9-DNA complex with all prominent peaks matching the peptides of our protein.

The mass spectrometric data of the Halo-ZnF^Cst^ 1-11 showed that there were no additional bacterial peptides in the complex based on searches of the NCBI or SwissProt databases. Next, the measured monoisotopic m/z value of the peptide mass fingerprint spectra was compared to the theoretical m/z values of Halo-ZnF^Cst^ 1-11 using ProteinProspector (University of California, www.proteinprospector.ucsf.edu). The 34 expected peptide ions from the expressed protein were detected in the peptide mass fingerprint spectrum (Table S2). This mass list was subjected to MS-Fit provided by ProteinProspector. The software tool MS-Fit could assign 18 m/z values to peptides correlated to Halo-ZnF^Cst^ 1-11, another 7 m/z values were identified manually. A sequence coverage of 59.58% was obtained. Four peptides resulting from autodigestion of trypsin and one CHCA-cluster were also identified manually. Four m/z values could not be assigned, but it can be assumed that these correspond to nonspecific fragments of Halo-ZnF^Cst^ 1-11.

To further confirm peptide mass fingerprint data, the MS/MS spectra of four prominent m/z values (1338.61, 1767.84, 1810.76, and 1908.01) were obtained (Figure S4B, Table S3). Analysis of these spectra was done by comparing measured m/z values to calculated values of the corresponding amino acid sequences: m/z 1338.61 SFIASEISSIER, m/z 1767.84 HQRTHTGEKPYVCR, m/z 1810.76 SDKPDLGYFFDDHVR, m/z 1908.01 LLFWGTPGVLIPPAEAAR. The amino acid sequences of all four peptides could be verified to be part of the protein construct. Persistent y-ion series in all four MS/MS spectra were detected, as well as, matching b and a ions. Mass lists showing matched peptide mass fingerprint and MS/MS data are included in the Table S2 and Table S3?.

## DISCUSSION

### PRDM9 binds DNA as a trimer

In this work, we developed an approach to infer the MW of the PRDM9-DNA complex from native gel electrophoresis using an *in vitro* binding assay (EMSA) with two independent strategies (*assay I* and *assay II*; Figure 2), which differed in the type of regression standards (lower-shift vs. free DNA, respectively) and inferred measurements (super-shift vs. lower-shift, respectively). Here we report that the multimer is formed by three PRDM9 units when actively bound to a specific DNA target sequence. It is possible that the ZnF array also forms larger or smaller complexes, but our gel images indicate that the majority of the active PRDM9 that specifically binds to DNA is formed by a trimer. This is congruent with previous studies also reporting that PRDM9 forms functional multimeric complexes of at least two or more units^14, 36^. Moreover, our data demonstrate that multimerization is independent of the tested PRDM9 allele (PRDM9^Cst^ and PRDM9^Dom2^). Neither functional tags like eYFP, MBP and Halo, nor expression systems (bacterial or *in vitro* expression) or protein purity did influence the binding or protein stoichiometry in all eleven tested protein constructs, confirming that our approach is very robust. By fingerprinting the PRDM9-DNA complex with mass spectrometry, we confirmed that the complex is solely formed by PRDM9 and DNA *in vitro.*

### PRDM9 multimerization is mediated within the ZnF domain

We removed the KRAB, SSXXD, PR/SET domain, the single ZnF0, and even shortened stepwise the PRDM9 ZnF array to the smallest construct with only five out of 11 ZnFs. In none of these constructs we observed a change in stoichiometry. Thus, we conclude that the PRDM9 multimerization is mediated within the variable DNA-binding ZnF domain. This is also congruent with a recent study using co-IP experiments of different co-expressed PRDM9 constructs reporting that PRDM9-PRDM9 interactions occur within the ZnF domain^14^. Moreover, five out of eleven fingers within the PRDM9^Cst^ ZnF domain, more precisely ZnFs 2-6, are sufficient for the formation of a stable trimer that binds specifically to DNA.

C2H2-type ZnF proteins, which form one of the largest groups of proteins identified so far, play a crucial role in many cellular processes like development, differentiation and tumor suppression (reviewed in^52^). There are three main types of C2H2 ZnF proteins, which are able to bind DNA sequences. These include the triple-fingered ZnFs (tC2H2) consisting of three consecutive fingers, like Zif268 or SP1, separated-paired ZnFs (spC2H2) with two fingers each grouped in pairs and separated from other pairs (like Tramtrack or Basonuclin), and multiple-adjacent ZnFs (maC2H2) having four or more fingers located closely in a row, like TFIIIA, Ikaros or Roaz (reviewed in^52^).

PRDM9 has many parallels to the maC2H2 subfamily. First, PRDM9 has an array of several ZnFs from 8 to over 20 fingers (reviewed in^18^). In comparison, TFIIIA is a transcription factor with nine adjacent C2H2 ZnF repeats, which binds to the 5S RNA promoter in *Xenopus laevis* oocytes^53, 54^. It contacts the DNA with fingers 1-3, but can also touch DNA at finger 5 and weakly binds to the DNA at fingers 7-9^55–58^. Another example is Zac, a seven C2H2 ZnF protein, promoting apoptosis and cell cycle arrest^59, 60^. It was shown that Zac has transcription activities upon binding DNA via the fingers 2-3 and 5-7, without including ZnF4^61^.

Only a subset of about 24-75% of maC2H2 ZnFs are part of the DNA sequence recognition; whereas the rest is free for other roles like RNA or protein-protein interactions (reviewed in^52^). Interestingly, those ZnFs of the maC2H2 family, which are not participating in DNA binding, often mediate dimerization, which can also increase the binding affinity, as it was observed for Ikaros^62^ and Roaz^63^. Similar to the other maC2H2 members, not all the ZnFs within the array of PRDM9 are necessary to form a stable and sequence-specific binding with DNA^12, 14^. Moreover, for the human PRDM9^A^ it was suggested, based on enrichment of DNA binding motifs, that ZnFs 5 and 6 within the array might interact weakly with DNA and instead act as linkers between up- and downstream ZnFs or might have other functions like a ZnF-ZnF interaction^14^.

In previous studies using yeast two-hybrid systems, co-transfection of isoforms and gel shift assays, complexes formed by maC2H2 (like Ikaros or Roaz) were interpreted as ‘higher order structures’ or dimers, but no exact stoichiometry was established^62, 63^. These ZnF proteins preferentially formed multimers in order to bind specific target DNA sequences with a higher binding affinity and efficiency (reviewed in^52^). Multimerization is usually mediated via ZnFs not participating in DNA recognition using two different modes: hydrophobic interactions through the ZnF surface^64^, as it was shown for proteins like GL1^65^ and SW15^66^, or ZnF-ZnF interaction mediated by the same amino acids conferring the DNA sequence specificity^67, 68^. Among others, this was shown for Ikaros, a hematopoietic cell-specific protein playing a major role in regulating lymphocyte development (reviewed in^69^). Ikaros consists of four adjacent ZnFs close to the N-terminus and are involved in sequence-specific DNA binding^70^; whereas, the two C-terminal ZnFs are highly selective for dimerization^67^.

In contrast, our truncation product ZnF2-6 of PRDM9^Cst^ showed that five ZnFs are sufficient to form a multimer. It is very likely that all these ZnFs are in direct contact with DNA, since at least five ZnFs are required for a stable and sequence-specific DNA binding, as was shown in several instances^12, 14^. Thus, in case of PRDM9 it seems more likely that a hydrophobic interaction of the ZnF surface confers the protein-protein interaction; however, we cannot exclude the possibility that in a longer PRDM9 ZnF array, a more specific ZnF-ZnF interaction might take place.

### Multimerization of PRDM9 is not exclusive of heteromers

ZnF proteins can form both homo- and heterodimers; however, they prefer interacting with the same protein rather than forming heteromeric complexes (reviewed in^52, 67^. So far, the question whether PRDM9 can form homo- and heteromeric complex has not been fully addressed. Baker and colleagues first discovered PRDM9 to multimerize and demonstrated that it can form active homo- and heteromeric complexes. This was shown in cells co-transfected with two identical PRDM9 allele constructs (PRDM9^C^) harbouring different tags (FLAG and V5) or with two different alleles, PRDM9^A^ and PRDM9^C^. Immunoprecipitation experiments using anti-FLAG followed by Western blot with anit-V5 showed that V5-PRDM9^C^ was only detected when co-expressed with FLAG-PRDM9^A^ or FLAG-PRDM9^C^. This was confirmed by Chromatin-Immunoprecipitation (ChIP) measuring the presence of PRDM9^A^ at C-defined hotspots when expressed simultaneously with PRDM9^C^. Moreover, when co-expressed with a catalytically-dead PRDM9^C^ mutant, PRDM9^A^ could replace the catalytic activity and trimethylated H3K4 at C-defined hotspots^36^.

Similar experiments expressing both human PRDM9^B^ and chimp PRDM9 suggested a higher preference of homo-than heteromeric complex formation. This was shown by competitive co-IP experiments where three constructs were co-transfected to cells harbouring different tags like chimp-V5, chimp-HA and human-HA. In this case chimp-HA was detected more efficiently than human-HA after IP pulldown for chimp-V5 and vice versa^14^. Similarly, no evidence for heteromer formation was observed *in vivo* by the trans-complement methyltransferase activity in mice heterozygous for PRDM9^Dom2^ and PRDM9^Cst-YF^ (PRDM9^Cst^ variant with a methyltransferase knockout mutation)^71^. The question is whether potential formation of a heteromer also depends on ZnF divergence, since those amino acids defining the variability do not only recognize specific DNA but also possibly mediate ZnF-ZnF interactions^67, 68^.

### The PRDM9 trimer binds only one DNA target

Since we showed that multimer formation is coordinated within the DNA-binding ZnF domain, we proposed different models of how many DNA molecules can be bound by the polymeric complex. Our data showed that the tandem-Hlx1 with two binding sites forms two independent complexes and that the PRDM9 trimer binds only to one DNA molecule in an equimolar mixture of long and short DNA sequences. This strongly suggests that the multimer only binds one DNA target molecule at a time, even though three ZnF domains would be available. It is possible that the two other ZnF domains are important in mediating stable ZnF-ZnF interaction, explaining why ZnF proteins are often found in multimers. However, we cannot exclude that the other domains of PRDM9 (e.g. KRAB, SSXRD or PR/SET) engage independently within the multimer. Previous reports have documented chromatin modification of H3K4me3 and H3K36me3 marks flanking meiotic recombination hotspots. The periodic methylation of these nucleosome marks decreases (in an asymmetric or symmetric fashion) with distance to the PRDM9 binding site^8,16,72,73^ (and reviewed in^18^). It is possible that the strong methylation signals located 2–3 nucleosomes up- and downstream of the PRDM9 binding site are the result of three active SET domains.

### What is the biological effect of PRDM9 multimerization

Given that the trimeric PRDM9 complex recognizes only one target molecule at a time, the effective dosage of PRDM9 may be affected especially in heterozygous individuals. It has been shown in several instances that PRDM9 dosage plays a crucial role in fertility with both homo- and hemizygous *Prdm9* null mice showing complete or partial sterility due to a drastically reduced number of active hotspots^7, 41^.

One explanation for *Prdm9* allele incompatibility in heterozygous individuals is the combination of both native and virgin PRDM9 target sequences of two evolutionary distant alleles within a certain genetic background, whereas mainly the virgin sequences are activated, resulting in hotspot asymmetry^33, 40^. Hotspot (a)symmetry may also be affected by the dominance of certain *Prdm9* alleles over others. As an example, it has been shown in humans and mice heterozygous for *Prdm9,* that hotspots specific for one allele have been enriched, suggesting that this allele is dominant, like PRDM9^C^ is dominant over PRDM9^A^ and PRDM9^Cst^ over PRDM9^Dom2^^5,33,41,74^. This may also depend on the binding affinity of PRDM9 for its target, which probably differs between variants^15^ and therefore affects PRDM9 dominance. In case of PRDM9 multimerization within a heterozygous context, different variants are physically coupled in a 2:1 ratio. This could affect the dosage of PRDM9, since one allele is over-the other is underrepresented, as well as, the distribution of (a)symmetric hotspots if one allele is dominant and binds better to its target. In this case, the weaker allele is outnumbered and probably even masked within the multimeric complex with its activity strongly suppressed within the multimer, leading to a more symmetric distribution of hotspots and thus higher fertility.

### Conclusions

Taken together, here we demonstrate an electrophoresis-based approach to investigate PRDM9 multimerization and observed a protein stoichiometry of a trimer. This trimerization is mediated within the highly variable DNA-binding ZnF domain, which only binds one specific target molecule at a time. With the possibility that two PRDM9 variants form a trimer in a 2:1 ratio within a heterozygous organism, we provide important insights in the nature of the ZnF-DNA interaction and DNA targeting of PRDM9 in general, but also in the context of hybrid sterility since dominance and dosage strongly correlate with both, PRDM9 homo-and heteromerization and sterility.

## MATERIALS and METHODS

### DNA sources

DNA fragments were produced via PCR amplification of genomic DNA of the B6 mouse using biotinylated or unmodified primers or hybridization of complementary single-stranded oligonucleotides. Details are shown in Supplementary_Methods.

### Cloning & expression of PRDM9 constructs

Distinct coding sequences of *Prdm9^Cst^* (CAST/EiJ strain, *Mus musculus castaneus* origin) and *Prdm9^Dom2^* (C57BL/6J strain, *Mus musculus domesticus* origin) were cloned into different vector systems for bacterial (pOPIN vector) and cell free *in vitro* (pT7-IRES-MycN vector) expression as it was described in Striedner et al.^12^. Therefore, the inserts were prepared via specific PCR amplification and cloned into the desired vector using restriction enzyme-based cloning. The different constructs were designed to involve different tags like His-tag, maltose binding protein (MBP) or enhanced yellow fluorescent protein (eYFP) as well as different parts of the *Prdm9^Cst^* or *Prdm9^Dom2^* coding region. One PRDM9^Cst^ construct included a His_6_-HaloTag (Halo), which was kindly provided by the Petkov Lab (Center for Genome Dynamics, the Jackson Laboratory, Bar Harbor, ME 04609, USA), used for bacterial expression. For the majority of expressed proteins, a crude lysate was used for further experiments. Only Halo-PRDM9^Cst^ ZnF1-11 was semi-purified by ion exchange chromatography based on a protocol described in Walker et al.^13^. A detailed description about cloning, expression, lysate preparation and purification can be found in Supplementary_Methods. In summary, we used the following constructs:

Construct 1: His-eYFP-PRDM9^Cst^ in pT7-IRES-MycN vector (*in vitro* expression system)
Construct 2: His-MBP-eYFP-PRDM9^Dom2^ ZnF in pOPIN-M vector (bacterial expression)
Construct 3: His-MBP-eYFP-PRDM9^Cst^ ZnF in pOPIN-M vector (bacterial expression)
Construct 4: His-eYFP-PRDM9^Cst^ ZnF in pT7-IRES-MycN vector (*in vitro* expression system)
Construct 5: His-PRDM9^Cst^ ZnF in pT7-IRES-MycN vector (*in vitro* expression system)
Construct 6: His-Halo-PRDM9^Cst^ ZnF1-11 in pH6HTN-His_6_-HaloTag-T7 vector (bacterial expression)
Construct 7: His-eYFP-PRDM9^Cst^ ZnF1-11 in pOPIN vector self-made (bacterial expression)
Construct 8: His-PRDM9^Cst^ ZnF1-11 in pOPIN vector self-made (bacterial expression)
Construct 9: His-eYFP-PRDM9^Cst^ ZnF2-11 in pOPIN vector self-made (bacterial expression)
Construct 10: His-PRDM9^Cst^ ZnF2-8 in pOPIN vector self-made (bacterial expression)
Construct 11: His-PRDM9^Cst^ ZnF2-6 in pOPIN vector self-made (bacterial expression)
Construct 12: His-MBP-PRDM9^Cst^ ZnF in pOPIN-M vector (bacterial expression) – not used for multimer assay

### Electrophoretic mobility shift assays (EMSAs)

#### General EMSA protocol

Different EMSA experiments did vary in terms of binding reactions, incubation and electrophoresis times but followed the general EMSA protocol described in Striedner et al.^12^. All details about EMSA experiments can be found in Supplementary_Methods.

#### Image analysis

Image analysis was performed using the Image Lab software 5.1.1 (Bio-Rad). The lanes and bands were defined manually, then the migration distances and pixel intensities could be quantified and analysed further using Excel and OriginPro software (Origin Lab).

### Inference of molecular weight of the PRDM9-DNA complex from native gel electrophoresis

We analysed two different PRDM9 alleles (PRDM9^Cst^, from the CAST/EiJ strain of *Mus musculus castaneus* origin; and PRDM9^Dom2^ from the C57BL/6J strain of *Mus musculus domesticus* origin) targeting specifically the DNA of the *Hlx1* or the *Pbx1* hotspot, respectively^9^. The PRDM9 protein was produced by bacterial or cell-free *in vitro* recombinant expression of different constructs carrying different tags, such as enhanced yellow fluorescent protein (eYFP), maltose binding protein (MBP), His_6_-HaloTag (Halo) or no tag. In addition, some of the domains of PRDM9 or repeats of the ZnF array were removed. In total, we tested eleven different protein constructs (for details see Supplementary_Methods): eYFP-PRDM9^Cst^, MBP-eYFP-ZnF^Cst^, eYFP-ZnF^Cst^, ZnF^Cst^, Halo-ZnF^Cst^ 1-11, eYFP-ZnF^Cst^ 1-11, ZnF^Cst^ 1-11, eYFP-ZnF^Cst^ 2-11, ZnF^Cst^ 2-8, ZnF^Cst^ 2-6 and MBP-eYFP-ZnF^Dom2^. This large range of different sized protein constructs varied in conformation and charge; yet, rendered similar relative mobilities in EMSA confirming that in our set-up the migration of the complexes was mainly dependent of its molecular weight.

#### Assay I

For multimer assay I we used the advantage of the tandem-Hlx1 molecules resulting in super-shift bands representing a second PRDM9 complex bound. Each experiment was used to analyse only one type of PRDM9 construct. The protein was bound to six single-Hlx1 (75bp, 740bp, 856bp, 1053bp, 1147bp, 1460bp) and four tandem-Hlx1 (114bp, 232bp, 352bp, 468bp), or three single-Hlx1 (75bp, 543bp, 740bp) and two tandem-Hlx1 (114bp, 232bp; for very small protein constructs, see Figure S2B) which increased in unspecific flanking sites. Protein-DNA binding complexes were separated by the sieving effect of a native 5% polyacrylamide gel driven by the negative charges of the DNA resulting in lower-shift (only one PRDM9 protein bound) or lower- and super-shift (one or two PRDM9 proteins bound, respectively) bands. A long (4368bp, usDNA1) and short (220bp, usDNA2) unspecific reference DNA were included, tested not to interact with the protein, which were then used to normalize for the migration distance of the different bands in each lane: ([usDNA2] – [usDNA1]) – ([lower-shift] – [usDNA1]) = [lower-shift]norm. The relative increase in [lower-shift]_norm_ compared to lane 1 was plotted against the relative increase in molecular weight [dMW]_lowershift_, which is given by the size of the DNA fragment, in a logarithmic scale resulting in a linear regression. [super-shift]_norm_ was then used to determine [dMW]_supershift_ based on the regression function. [dMW]_supershift_ – [dMW]_lowershift_ = [dMW] for each tandem-DNA sample represents one additional PRDM9 complex. By using the molecular weight of the monomeric PRDM9 construct (e.g. 55kDa for ZnF^Cst^), the protein stoichiometry (#PRDM9) can be calculated from [dMW]. With four tandem-Hlx1 DNA fragments, four values for protein stoichiometry have been observed for each experiment. The experiments for one type of PRDM9 construct were replicated at least three times.

#### Assay II

In order to evaluate the *multimer assay II* experiments, one EMSA was used to investigate eight different types of PRDM9 constructs which was replicated for four times. All protein constructs were bound to a DNA fragment of 75bp (for PRDM9^Cst^ constructs the Hlx1 hotspot was used; for PRDM9^Dom2^ construct the Pbx1 hotspot was used). Unspecific reference DNA fragments of 2585bp (usDNA1) and 75bp (usDNA2) were included in each lane. The calculation of the PRDM9 stoichiometry was performed the same way as for *assay I.* However, a ladder of unbound DNA (75bp, 114bp, 273bp, 543bp, 740bp) were used as standards instead of the lower-shift. The stoichiometry was derived from the migration distance of the lower-shift band.

More details to calculate the protein stoichiometry using multimer *assay I* and *assay II* can be found in Supplementary files in Table S7 and S8.

### Statistical analysis

We tested for significant differences of calculated protein stoichiometry between different PRDM9 constructs for *assay I and II* separately using an ANOVA taking normality and homoscedasticity into account. A detailed description of the statistical analysis can be found in Supplementary_Statistical_Analysis.

### Mass Spectrometry

#### Chemicals

Acetone p.a., acetonitrile p.a. (ACN), acetic acid 96%, ethanol 96% (EtOH) were obtained from Merck (Darmstadt, Germany). Alpha-cyano-4-hydroxycinnamic acid (CHCA), ammonium hydrogen carbonate (NH_4_HCO_3_), coomassie brilliant blue R250 (CBB), dithiodreitol (DTT), iodoacetamide (IAA), trifluoroacetic acid (TFA) were obtained from Sigma-Aldrich (St. Louis, Missouri, USA). 5% mini-PROTEAN TBE gel was obtained from BioRad (Munich, Germany). Sequencing grade modified trypsin was obtained from Promega (Madison, Wisconsin, USA) und C_18_ ZipTips from Merck Millipore (Burlington, Massachusetts, USA).

#### DNA preparation

A single-stranded DNA fragment was extended to produce the 75bp fragment of the murine *Hlx1* hotspot. Therefore, 25μM of the synthetic oligonucleotide ssHlx1-75b was hybridized with 25μM of the primer single-Hlx1_R1 (sequences are listed in Table S4) in a 30μl reaction by incubating for 5 minutes at 95°C and cooling down for one hour. The hybridized DNA sample was supplemented with 1x NEB buffer 2.1 (NEB), 1mM dNTPs (Biozym) and 6.75 units T4 DNA polymerase (NEB) in a 56μl reaction and incubated for 1 hour at 12°C to start DNA extension. To remove remaining single-stranded DNA fragments, the sample was digested with Exonuclease I (NEB) as described in Supplementary_Methods. In order to purify the DNA, the sample was mixed with 2μl Co-Precipitant Pink (VWR) and 0,5 volumes of 5M NH_4_OAc. Furthermore, two volumes of pure ethanol were added and mixed by inverting. For total DNA precipitation, the sample was incubated at −20°C for 30 minutes followed by centrifugation at maximum speed for 30min at 4°C. The supernatant was discarded and the pellet washed with 1ml 80% ethanol. After a final centrifugation step of 5min at full speed and 4°C, the supernatant was carefully discarded and the pellet was dried at room temperature. The DNA sample was dissolved in 20μl nuclease-free water (Sigma-Aldrich).

#### Prepare binding reaction

In order to prepare the PRDM9-DNA binding complex, 7μl of semi-pure Halo-ZnF^Cst^ 1-11 were mixed with 2μM Hlx1-75bp DNA in a 20μl binding reaction supplemented by 1x binding buffer (10mM Tris, 50mM KCl, 0.05% NP-40, 50μM ZnCl_2_) and incubated for 60min at room temperature. The reaction was prepared twice.

#### Gel electrophoresis

20μL sample solution was supplemented by 1x DNA loading dye (Thermo Scientific) and applied onto the gel. Electrophoresis was performed on 5% mini-PROTEAN TBE (Biorad), 10 wells 30μL gels using 1xTBE (89mM Tris, 89mM boric acid, 3mM EDTA) as running buffer. Constant voltage was set to 100V (50mA/gel), after 40 min the separation was stopped.

After gel electrophoresis coomassie staining with CBB R250 was performed. The gel was fixed (45% EtOH, 5% acetic acid in water) for 45min and subsequently stained (0,1% CBB R250 in 45% EtOH, 5% acetic acid in water) for one hour. Destaining was performed using two solutions: destain solution I (40% EtOH, 7% acetic acid in water) for 30min, followed by destain solution II (5% EtOH, 7% acetic acid in water) overnight for clearing the background to obtain distinct protein bands.

#### In-gel tryptic digestion

The protein gel band was excised and cut into small cubes. To remove contaminants and CBB R250 stain various washing steps each lasting 15-min were applied: once with water, two times with ACN/water (1:1), once with 100% ACN and once with ACN/50mM NH_4_HCO_3_ pH8.5 (1:1, v/v). Gel pieces were dried in a vacuum centrifuge. Subsequently disulfide bridges were reduced with 100mM DTT (15.4 mg/mL in 50mM NH_4_HCO_3_ pH 8.5) for 45min at 56°C and alkylated with 55mM iodoacetamide (10.2 mg/mL in 50mM NH_4_HCO_3_ pH 8.5) for 30min at room temperature in the dark. Another washing step with ACN/50mM NH_4_HCO_3_ pH 8.5 (1:1) was performed. Gel pieces were dried in the vacuum centrifuge. Subsequently the gel pieces were incubated with 15μL digestion solution (12.5ng/μL Trypsin in 50mM NH_4_HCO_3_) for 15min and then coated with 25μL 50mM NH_4_HCO_3_ pH 8.5. The protein was digested at 37°C overnight.

Peptide extraction from the gel pieces was performed by using ACN/50mM NH_4_HCO_3_ pH 8.5 (1:1), ACN/0.1% TFA (1:1) and 100% ACN each step lasting 15min. Extracts were pooled and lyophilised in a vacuum centrifuge.

#### MALDI sample preparation

First, the stainless steel MALDI target was prepared by application of 1μL CHCA matrix solution (6mg/mL in acetone). After evaporation of acetone at room temperature a thin homogenous layer of matrix crystals was obtained.

Peptides were dissolved in 0.1% TFA and desalted using C_18_ ZipTips. The tips were activated with ACN/0.1% TFA (1:1) and equilibrated with 0.1% TFA. After binding of the peptides, salts and detergents were removed by washing the tips five times with 0.1% TFA. Elution was performed using 1.5μL ACN/0.1% TFA (6:4) which were directly applied onto the prepared CHCA layer on the MALDI target. The sample spot was dried at room temperature and subsequently transferred into the AXIMA Performance instrument.

#### Instrumentation

Gel electrophoresis was performed on a Mini-PROTEAN (BioRad) vertical electrophoresis cell connected to a Consort EV265 (VWR; Radnor, Pennsylvania, USA). MALDI-TOF spectra were acquired on an AXIMA Performance instrument (Shimadzu; Kyoto, Japan). The AXIMA Performance is equipped with a nitrogen laser (λ=337nm) and it was operated in positive ion, reflectron mode using pulsed extraction. Peptide mass fingerprint (PMF) mass spectra were acquired by averaging 500 and MS/MS spectra by averaging up to 2500 unselected and consecutive laser shots. No smoothing algorithm was applied prior to data analysis.

## Acknowledgements

This work was supported by the ‘Austrian Science Fund’ (FWF) 27698-B22 to I.T.-B. Open access funding was provided by Johannes Kepler University Linz. We are grateful to Petko Petkov for providing the vector containing the coding sequence of the *Prdm9* alleles of the strains CAST/EiJ and C57BL/6J, the mouse genomic DNA of these two strains and the vector containing the Halo-ZnF^Cst^ 1-11 construct. We also want to thank Hermann Gruber for fruitful discussions and Angelika Heißl for comments and input in the project.

## Author contributions

I.T.-B. and T.S. conceived the research; T.S., Y.S., J.K. and I.T.-B. designed the experiments; T.S., Y.S., K.D., J.K. and N.Z. performed the experiments; I.T.-B. contributed new reagents/analytic tools; T.S., Y.S., P.H. analysed the data; and T.S., Y.S., J.K. and I.T.-B. wrote the paper. All authors read and approved the final manuscript.

## SUPPLEMENTARY FILES

Supplementary notes, supplementary figures, supplementary tables, supplementary methods, and supplementary statistical analysis can be found in the supplementary files.

